# Short-term radiofrequency exposure from new generation mobile phones reduces EEG alpha power with no effects on cognitive performance

**DOI:** 10.1101/363929

**Authors:** Zsuzsanna Vecsei, Balázs Knakker, Péter Juhász, György Thuróczy, Attila Trunk, István Hernádi

## Abstract

Although mobile phone (MP) use has been steadily increasing in the last decades and similar positive trends are expected for the near future, systematic investigations on neurophysiological and cognitive effects caused by recently developed technological standards for MPs are scarcely available. Here, we investigated the effects of radiofrequency (RF) fields emitted by new-generation mobile technologies, specifically, Universal Mobile Telecommunications System (UMTS) and Long-Term Evolution (LTE), on intrinsic scalp EEG activity in the alpha band (8–12 Hz) and cognitive performance in the Stroop test. The study involved 60 healthy, young-adult university students (34 for UMTS and 26 for LTE) with double-blind administration of Real and Sham exposure in separate sessions. EEG was recorded before, during and after RF exposure, and Stroop performance was assessed before and after EEG recording. Both RF exposure types caused a notable decrease in the alpha power over the whole scalp that persisted even after the cessation of the exposure, whereas no effects were found on any aspects of performance in the Stroop test. The results imply that the brain networks underlying global alpha oscillations might require minor reconfiguration to adapt to the local biophysical changes caused by focal RF exposure mimicking MP use.

## Introduction

The worldwide use of mobile phones (MPs) is still rapidly growing. By 2020, almost three-quarters of the world’s population—or 5.7 billion people—will subscribe to mobile services.^1^ When MPs are used in close proximity to the head, it absorbs a considerable portion of the energy of radiofrequency (RF) output power.^2^ To date, research does not offer any consistent evidence concerning the adverse health effects from short-term exposure to RF fields at levels below those that cause tissue heating.^3,4^ However, given the large number of MP users, it is important to investigate, understand and monitor any potential public health impact^5^ they may pose. Owing to this, there has been an increasing concern about the effects of RF exposure emitted by MPs on psychophysiological and cognitive indices of brain function.

Third-generation (3G) MPs use Wideband Code Division Multiple Access (WCDMA) standard RF signals, also called Universal Mobile Telecommunication System (UMTS), which operate in the frequency range of 1920 to 2170 MHz. The WCDMA modulation is a non-periodic mixture of signals with 5-MHz bandwidth.^6^ Fourth-generation (4G) mobile services are based on the Long-Term Evolution (LTE) technology, which is the recent generation global mobile communication standard. The LTE service may utilise all of the frequency bands currently in use by legacy communication technologies (e.g. 1710–1785 MHz (uplink) and 1805–1880 MHz (downlink)), as well as a few new bands (e.g. 2500–2570 MHz (uplink) and 2620–2690 MHz (downlink)). LTE is a fully digital, Internet Protocol-based technology. In 2016, new-generation connections (3G and 4G technologies) accounted for 55% of total mobile broadband connections, and the proportion of 4G connections alone is forecasted to reach 41% by the end of the decade.^1^

Previous papers focused mainly on health effects of exposure to RF emitted by the classic second-generation (2G) MPs using the Global System of Mobile (GSM) technology.^7,8^ In contrast, only a few studies examined the possible effects of 3G mobile radiation on cognition,^9-15^ whereas for the LTE system used by 4G MPs, no indices of human cognitive performance have been investigated thus far. Moreover, findings regarding the possible cognitive effects of MP use are rather inconsistent; for example, some suggest adverse^11,16,17^ or facilitating effects,^18-21^ whereas others found no effects.^12,14,22-26^ Previous reviews do not support the short-term impact of high frequency electromagnetic fields (EMF) emitted by several types of MPs on human cognitive performance (for an overview, see the following reviews: ^7,8,22,27,28^). It is argued that the heterogeneity of the results may be due to differences in methodology, statistical power and interpretation criteria.^27^ Nevertheless, almost all cited studies examined the effects of the classic GSM system, with only a limited range of cognitive tasks applied.^28^ Most studies applied simple reaction time (RT) measurements or go/no-go tests.^27^ Based on a recent review study, GSM-like low-intensity EMF exposure does not seem to cause any measurable cognitive and/or psychomotor effects measured in simple reaction tasks.^29^ However, despite the fact that evidence suggests that basic sensorimotor mechanisms probed in simple tasks are robust to EMF exposure, subtle disturbances may well be caused by RF EMF in more complex and thus more vulnerable cognitive processes, which should be investigated using suitable experimental paradigms. Given that the Stroop test measures selective attention, cognitive flexibility, processing speed and executive functions within the same task, it enables the testing of various higher-order cognitive functions well beyond the capacity of the usually applied simple RT tasks.^30,31^ Certain mental illnesses, including dementias,^32,33^ schizophrenia^34,35^ and attention-deficit hyperactivity disorder (ADHD)^36,37^ have been successfully screened by measuring the size of the so-called Stroop-effect (see below) within the task. Thus, by using the Stroop test, complex and finely tuned cognitive networks can be investigated.

The Stroop-effect reflects various cognitive phenomena, especially (semantic) interference, facilitation and the automatic nature of reading.^30^ *Stroop interference* (IF) occurs when the written word on the screen is incongruent with the colour of the font used to write that word (e.g. the word ‘green’ written in blue). Longer response times can be recorded in the incongruent condition relative to the neutral condition (in which the conflicting information is absent from the stimulus), as the subject has to suppress automatic reading to determine colour. *Stroop facilitation* (FAC) occurs in the congruent condition (e.g. the word ‘green’ written in green), as reading the matching word speeds up colour-naming, which is manifested in shortened response time. Both IF and FAC appear only in the colour-naming (CN) task condition and not in the word-naming (WN) task condition.^38^

The most prominent methods for non-invasive measurements of human brain function are magnetoencephalography (MEG), electroencephalography (EEG) and functional magnetic resonance imaging (fMRI). Most previous research on the effects of RF on human brain function utilise EEG, presumably because it has excellent temporal resolution, it is widely available (relatively to MEG) and the measurement itself does not involve the use of RF pulses (unlike fMRI). Although scalp-measured EEG is the result of spatiotemporal summation of a plethora of cellular level processes, decades of research suggest that it can still provide rich and valuable information of the neurophysiological processes underlying healthy and pathological brain function.^39-41^ A recent review article about the effects of EMF on resting EEG^42^ covered 36 experiments (minimum criteria for inclusion were randomised, single-blind studies with crossover procedure, and mostly involved GSM MPs) where MP-EMF effects were tested on resting wakeful EEG in healthy humans, 72% of which confirmed the existence of the EMF-EEG relationship, most prominently in the alpha (8–12 Hz) frequency band. Most of these studies involved second-generation (2G) MP technology, the methodology was very heterogeneous and most reviews concluded that the findings varied even in the direction of effects, and thus are hard to reconcile. In addition, a systematic investigation of UMTS and LTE technologies with identical EEG methodology has not been performed thus far, and the possible cognitive effects of exposure to LTE RF have remained unexplored.

In an attempt to contribute to the systematic investigation of RF exposure effects, we studied the neurophysiological and cognitive effects of UMTS and LTE technologies within the same experimental framework. The question we sought to answer was whether short-term (20 minutes) RF exposure with the two most widely used new-generation technologies (3G, 4G) would induce any measurable effects on brain function as indexed by intrinsic alpha band EEG activity and cognitive performance assessed by the Stroop test.

## Materials and Methods

### Participants

Two identical investigations were performed, only differing in the type of RF exposure used (UMTS and LTE), which are hereinafter referred to as UMTS experiment and LTE experiment. Thirty-four healthy participants (20 females, aged 20 ± 3years) participated in the UMTS experiment. In the LTE experiment, 26 healthy participants were involved (13 females, aged 21 ± 3 years).

Participants were right-handed and medication-free according to self-report. They were fulltime students of the University of Pécs, Hungary. All participants had normal or corrected-to-normal vision. Participants reported that they use their right hand for phone calls. They were instructed to abstain from smoking, alcohol and caffeine consumption at least six hours prior to the investigation and to moderate phone use before investigation (less than 30 minutes on investigation day). They were compensated by practical course credit for the time spent in the experiment. All participants signed a written informed consent after the experiment was fully explained. The study was conducted in accordance with the declaration of Helsinki and the protocol was approved by the Regional and Institutional Research Ethics Committee of the University of Pécs (file number 5348). Recordings were carried out in the Psychophysiology Laboratory of the Department of Experimental Neurobiology, University of Pécs.

### Experimental design

Two separate experimental sessions were conducted with at least a one-week interval between the sessions. The timelines of the two sessions are shown in Fig. 1. Both sessions were carried out at an identical time of the day (in the morning or in the afternoon) between 8 am and 6 pm, balanced across participants to avoid possible interference with circadian regulation effects.^12^ Participants were seated in a comfortable armchair in front of the screen in a dimly lit room. Both the subject and the experimenter were present in the recording room, but no visual contact was possible during the Stroop tests and EEG recordings. To sustain the alertness of subjects, a muted nature film was played during the EEG recording; the film was started exactly at the beginning and stopped at the end of the resting EEG period (Fig. 1). This controlled visual stimulus contributed to diverting the possible imagery of the investigation or of the RF itself. The order of the two documentary clips, the order of the RF exposure types (Real, Sham) across the two sessions, and the order of the Stroop tasks (WN, CN) of one individual Stroop test within one session and between sessions were randomly assigned and counterbalanced across participants. Exposure to RF energy was administered in a double-blind fashion.

**Figure 1.**
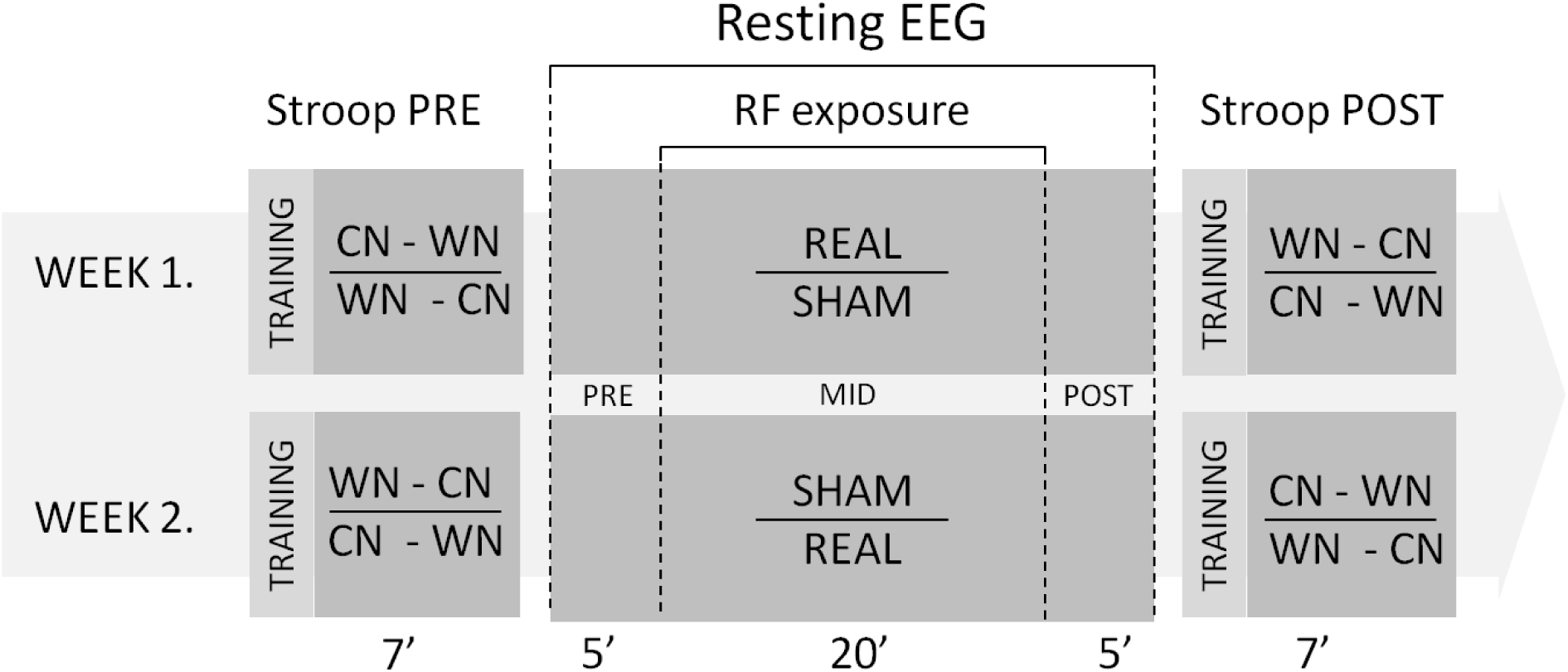
Schematic illustration of experimental design. First, the Stroop test was performed. After that, a 30-min long rEEG was registered continuously. After five minutes of EEG recording (Pre), RF exposure (either Real or Sham) was turned on and lasted for 20 minutes (Mid). Five additional minutes rEEG (Post) was registered continuously after ending of exposure. After that rEEG registration (and the documentary clip) was stopped, the RF exposure headset was quickly removed and participants did the second Stroop test of the session. The data of the two studies were collected by the same person, and later processed identically at the same time. The random elements of the counterbalancing are separated by horizontal line.

### Stroop test

A computerised version of the Stroop colour word test was designed to assess interference and facilitation effects between two different types of information: colour and word. The test was programmed in the freeware experiment programming language PEBL,^43^ a free psychology software package that allows users to create or develop experiments freely without licence or charge (http://pebl.sourceforge.net). Test stimuli (Fig. 2) were coloured words sequentially projected on the screen and the participants had to respond with appropriate key presses according to the task rules on a keyboard from which task-irrelevant buttons were removed. RTs were recorded in ms accuracy on a PC microcomputer for later processing. Stimuli were projected on a screen at a distance of approx. 1.4 m in front of the participant, subtending vertical visual angles of approx. 0.3°. The monitor refresh rate was 120 Hz and the response time was 2 ms. Every Stroop test consisted of three blocks: one training block and two task blocks: CN (in which one had to react to the colour of the word—response by shade) and WN (in which one had to establish the meaning of the word—response by meaning). Each block consisted of only one kind of task. The type of the initial task (WN, CN) was randomly selected for each participant and then alternated within and between sessions (see Fig. 1, mixed elements are separated by horizontal lines). The number of trials per block was 20 in the training block and 60 in the task block, with an equal number of trials from each condition. Stimuli were coloured words “red”, “green”, “blue”, “yellow” (in Hungarian). Three stimulus conditions were used: congruent (=, colour name and colour match), incongruent (≠, colour name does not match colour) and neutral (×, either the colour is black or colour name is ‘xxxxx’) as control stimuli for comparison purposes.

**Figure 2.**
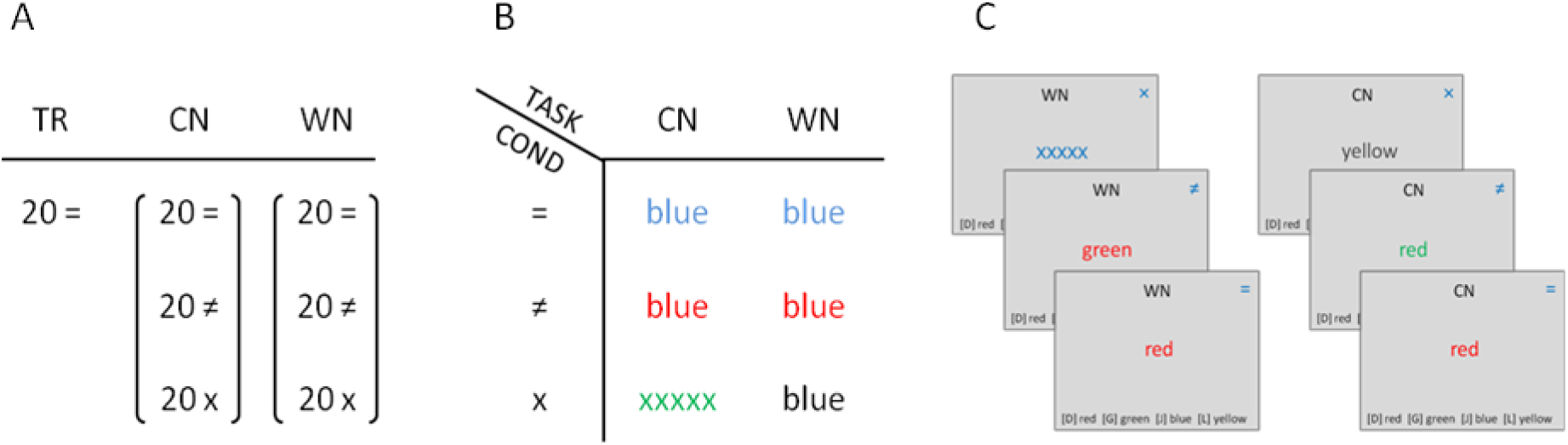
Arrangement of the Stroop test. Each individual Stroop test consisted of 140 stimuli: 20 training stimuli and 120 test stimuli in equal amounts per task types and per conditions. Abbreviations: [TR] training, [TASK] task – [CN] colour-naming: to pay attention to the colour of the font (ignoring the word itself), word-naming [WN]: to pay attention to the word itself (ignoring the colour of the word); [COND] condition—congruent [=]: colour and name of the word is the same; incongruent [≠]: colour and name differ; and neutral [×]: neither the colour nor the name is among the targets. (A) Number of stimuli per one Stroop test. (B) Stimulus types. (C) Schematic depictions of stimulus layout. The words here are in English, but in the study, we used words in the participants’ mother tongue (in Hungarian).

### Stroop test data analysis

To test differences in performance in the two separate sessions (Real/Sham exposure), RTs were recorded and median RT values of different task conditions were computed. Considering the median of RTs instead of mean stems from a concern that RT distributions are positively skewed—similar to the ex-Gaussian distribution^44,45^—and the median value is less affected by outliers and less sensitive to deviation from normality.^46^ Based on the distribution of all RTs from all participants in the Sham condition, responses with RTs more than two standard deviations above the mean were considered slow. Participants with more than 10% erroneous responses or more than 10% slow responses to non-training targets were excluded from data analysis, assuming that they were not sufficiently motivated or focused on the task.^47^ Only the RTs of correct responses of remaining participants were analysed. In the UMTS experiment, 7 of the 34 participants were excluded from data analysis as exclusion criteria were met: 5 participants were excluded due to errors and 2 due to extreme slowness. In the LTE experiment, 3 of the 26 participants were excluded: 2 participants due to errors and 1 due to extreme slowness.

RT data were analysed with mixed ANOVA to assess possible RF exposure effects on cognitive performance. As IF and FAC are surely detectable only in the CN task,^38^ RTs of the WN task were not the focus of statistical analysis. Interference was probed by (2 × 2 × 2) × 2 ANOVA (within-subject factors: exposure (two levels: Real, Sham exposure) × time (two levels: Pre, Post), IF (two levels: CN incongruent, CN neutral)), between-subject factor: RF type (UMTS, LTE). Likewise, FAC was tested with ANOVA (within-subject factors: exposure (two levels: Real, Sham exposure) × time (two levels: Pre, Post), FAC (two levels: CN congruent, CN neutral)), between-subject factor: RF type (UMTS, LTE). In those significant interactions where condition IF or FAC were concerned, post-hoc tests were further investigated. Alpha level was set to 0.05 and Tukey correction was used to control for multiple testing. Assumption checks of normality and of sphericity (Mauchly’s Test of Sphericity) were conducted.

Responses are fastest in the congruent condition because of facilitation and slowest in the incongruent condition due to interference, so the difference in RT between these conditions (congruent minus incongruent) yields a cumulative index of the two Stroop-effects (CUM), which is potentially more sensitive to concordant changes in the IF and FAC effects. CUM was studied with a design similar to that of the previous ones: (2 × 2 × 2) × 2 ANOVA (within-subject factors: exposure (two levels: Real, Sham exposure) × time (two levels: Pre, Post), CUM (two levels: CN incongruent, CN congruent)), between-subject factor: RF type (UMTS, LTE).

### Subjective ratings

At the end of the second session in the UMTS experiment, participants were asked the following question (in Hungarian): “What do you think, in which of the two sessions did you get exposed to the actual, Real irradiation? At the first (A) session? /At the second (B) session? /No idea?” Fourteen of them correctly identified the session of Real exposure, 14 participants gave a wrong answer and 6 could not tell the answer. When A and B responses in which the subjects expressed their uncertainty (saying e.g. “…but it is only a guess”) were counted as “No idea”, the proportion of right and wrong answers was still similar: 11 right, 10 wrong, 13 no idea. Thus, we can conclude that participants in the UMTS experiment really could not establish when the Real UMTS exposure was applied. In the previously conducted LTE experiment, no subjective ratings query was done.

### RF exposure

#### UMTS experiment

The UMTS MP exposure system was developed in our laboratories and was successfully used in previous studies.^14,48-51^ The UMTS RF source was a standard Nokia 6650 (Nokia, Espoo, Finland) MP operated by Phoenix Nokia Service Software (v. 2005/44_4_120; Nokia, Espoo, Finland). The MP was connected to an RF amplifier (Bonn Hungary Electronics Ltd., Hungary) via the external antenna output of the MP. A patch antenna (Reinheimer Elektronik, Wettenberg, Germany; model no: M30EXO-0250-XX) was connected to the output of the RF amplifier. The patch antenna was mounted in a position mimicking the normal use of an MP: the centre of the patch antenna was near the exit of the ear canal, above the tragus, at a distance of 7 mm. A two-position switch located on the front panel of the RF amplifier enabled double-blind experimental conditions: one position was associated with Real exposure, and the other with Sham exposure. Neither the investigator nor the participants were aware of the actual exposure condition. The system was set to WCDMA mode operating at the 1947 MHz carrier frequency (which corresponds to the operating frequency of UMTS MPs in Europe) with a wideband 5 MHz modulation of the RF carrier signal.^6^ SAR measurements were performed using the same exposure setup as in this study in Trunk *et al.* (2013).^49^ The averaged SAR was below 2 W/kg in any position within the phantom, meeting the limit of public exposure to RF radiation in the European Union (EU) Recommendation [1999/519/EC Recommendation, Brussels, Belgium]. The duration of exposure was 20 minutes. For a detailed description of exposure, see Supplementary Material 1 (Exposure Systems and RF Dosimetry).

#### LTE experiment

The LTE RF exposure system consisted of a programmable signal generator, a power amplifier, the antenna holding fixture and the same patch antenna as in the UMTS study. The LTE “signal cocktail” was generated by an Anritsu MG3700A (Anritsu Co., Japan) programmable signal generator. The signal generator was remotely controlled by a computer in a way that was not visible to the investigator or the participants, either enabling or disabling signal output (corresponding to Real or Sham exposure, respectively), thus enforcing double-blind experimental conditions. For this experiment, we chose a carrier frequency of 1750 MHz (corresponding to one of the operating frequencies of LTE systems in Europe), and the LTE signal used 20 MHz of bandwidth (the allowed maximum). The signal generator was connected to the RF power amplifier BPAM14 (Bonn Hungary Electronics Ltd., Hungary). In our experiment, we set the maximum peak SAR to 1.8 W/kg and the ear-antenna distance to 7 mm. Figure 3 shows a block diagram of both exposure setups. For a detailed description of exposure, see Supplementary Material 1 (Exposure Systems and RF Dosimetry).

**Figure 3.**
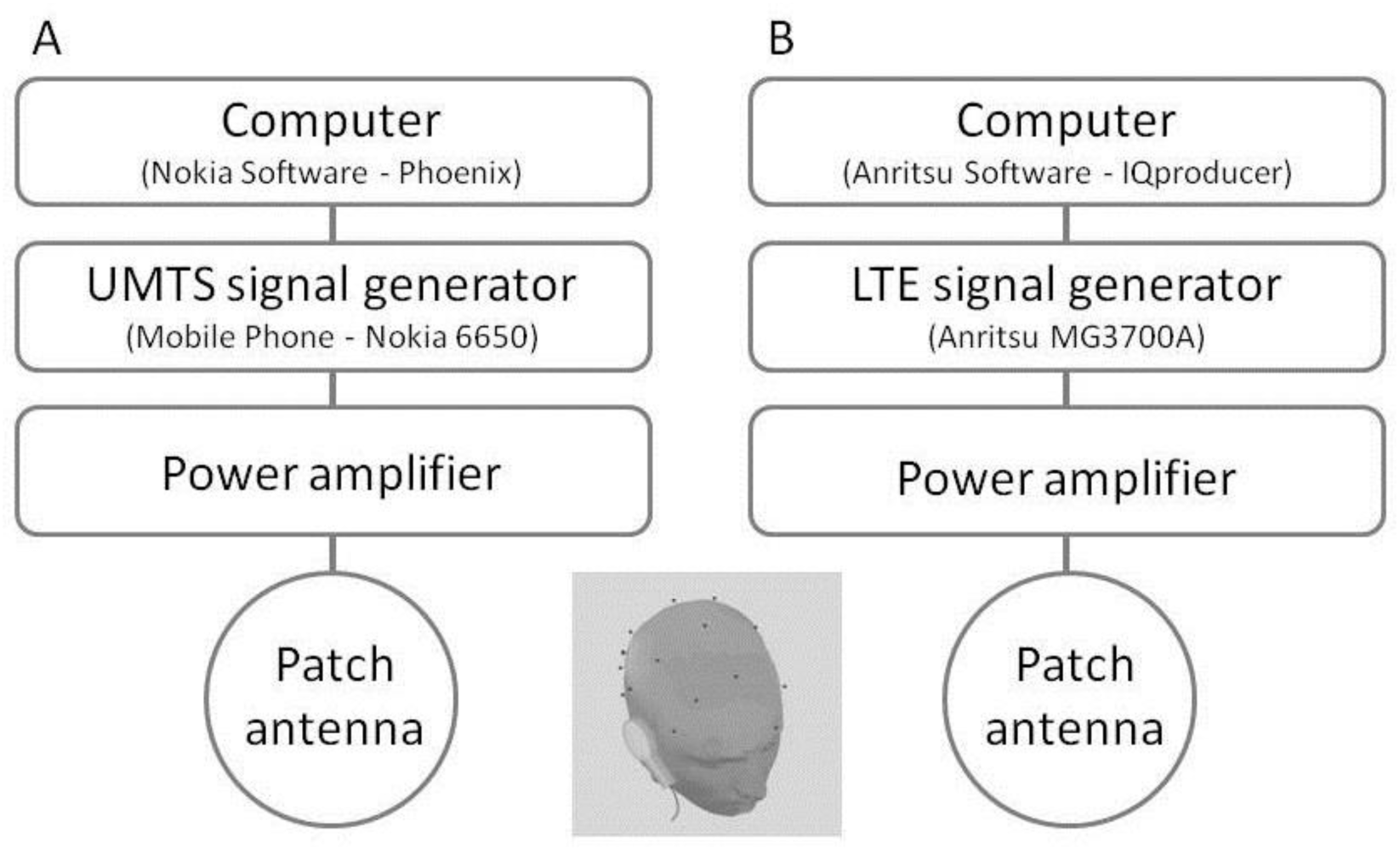
Scheme of UMTS (A) and LTE (B) exposure system.

### EEG data acquisition

Participants were fitted with an elastic EEG cap (Easycap, Munich, Germany) mounted with 32 Ag/AgCl electrodes positioned according to the international 10/20 System, referenced to the nose and grounded on the forehead. Data were recorded with a BrainAmp amplifier (Brain Products GmbH, München, Germany) at sampling rate of 1 kHz, band-pass filtered online with 0.1 Hz low and 450 Hz high cut-off and 45–50 Hz notch. Electrode impedances were below 5 kOhm at the beginning of each session. Electrooculogram (EOG) was recorded from the outer canthus of the right eye for off-line artefact rejection. During the recording, to minimise muscle-derived artefacts, participants were asked to avoid head and eye movements as much as possible. In each session, participants watched a documentary clip with the sound turned off during EEG recording. They were informed that no film-related tasks would be given during the experiment.

### EEG data processing

#### Pre-processing

EEG data were analysed off-line on a personal computer using EEGLab 14.0.0b^52^ in the MATLAB software environment (Mathworks Inc., Natick, MA, USA). Continuous EEG data were cut into three consecutive segments: block *Pre* (before exposure–5 min), block *Mid* (during exposure–20 min) and block *Post* (after exposure–5 min). EEG data from each block was filtered with zero-phase Hamming-windowed sinc filters (eegfiltnew.m in EEGLAB): a high-pass filter with a passband edge at 0.5 Hz (−6 dB cut-off at 0.25 Hz, filter order: 6601), a low-pass filter with a passband edge at 80 Hz (−6 dB cut-off at 95.625 Hz, filter order: 157) and a notch filter with passband edges at 45–55 Hz (−6 dB cut-offs at 46–54 Hz, filter order: 1651). Independent component analysis (ICA) was computed to decompose the EEG signals. ICA was only used to aid artefact rejection (see below), no components were removed from the data, and all the analyses were conducted in channel space. Data were segmented into 2 s epochs, yielding 150 Pre, 150 Post and 600 Mid segments. Bad epochs were rejected using a semi-automatic method based on ICA component time series. Bad epochs were first marked based on the joint probability distribution of all trials, kurtosis and temporal trends. The thresholds were as follows: 6 standard deviations for single-component kurtosis and probability, 5 standard deviations for all-component kurtosis and probability, a maximum slope (across the 2 s epochs) of 10 and an R^2^ limit of 0.8. After this, all segments were visually inspected to discard segments potentially missed by the automatic method. Less than 10% of epochs were rejected. Data were re-referenced to the average of all channels, and a fast Fourier transformation (FFT) was applied to artefact-free, epoched data with 1 Hz resolution to obtain spectral power values. Log-transformed power values were calculated for all electrodes within the alpha (8–12 Hz) frequency band for each block for statistical analysis.

#### Statistical analysis

First, alpha power averaged across all EEG channels (the EOG channel was excluded) was analysed with three-way (2 × 3 × 2) mixed ANOVA: within-subject factors: exposure (two levels: Real, Sham exposure) × time (three levels: Pre, Mid, Post), between-subject factor: RF type (UMTS, LTE). To further characterise exposure effects across time, a follow-up analysis was conducted where pre-exposure alpha power was subtracted from mid- and post-exposure values (hence, the time factor has only two levels here, Mid and Post). Further ANOVAs were performed to examine the effects of the two different RF types separately.

After this, we examined the topographic distribution of the effects using mass univariate analyses that were analogous with the ANOVA analyses we conducted. First, we tested for the exposure effect pooling data across RF types, followed by separate analyses for UMTS and LTE exposure. Finally, we explicitly tested whether exposure effects differed between UMTS and LTE exposure types. Each analysis was run on all three time windows and all the channels, yielding 90 tests per analysis (3 time windows × 30 electrodes); the within-analysis familywise error rate (α = 0.05, two-sided) was controlled by means of the cluster-based permutation method as implemented in the FieldTrip toolbox.^53,54^ The tests were based on parametric statistics, which were paired-sample t-tests for exposure effects and independent samples t-test on individual Real minus Sham difference scores calculated for each subject for the exposure × RF type interaction. Spatiotemporal clusters were formed using the conventional two-sided α_Cluster_ = 0.05 threshold of the local t-tests. During the clustering, channels were considered neighbours if they were closer than 5 cm from each other, except for Fp1, Fp2, O1 and O2, for which this threshold was 7 cm, because they were more distant from the rest of the electrodes. No minimal spatial cluster extent (in terms of minimum number of neighbouring channels) criterion was set. The number of permutations was 9999 and the maximum sum cluster statistic was used. The resulting p-values, reported as p_Cluster_, were multiplied by 2 to account for the fact that positive and negative effects were tested against separate null distributions (cfg.correcttail = ‘prob’ option in FieldTrip).

Although factors were carefully balanced, additional ANOVAs were conducted as control analyses involving factors that may have affected our results, the order of exposure sessions and the order of movies played in the two sessions. The two ANOVAs were designed as follows: within-subject factors: exposure (two levels: Real, Sham exposure) × time (three levels: Pre, Mid, Post), and one of them with between-subject factor session order (Real exposure on first week, Sham exposure on first week) and the other with the between-subject factor film order (film A on first week, film B on first week).

## Results

### The effect of exposure on Stroop-effects

During Stroop CN task performance, participants gave slower responses in the incongruent condition (IF: F_1,48_ = 132.155; p < 0.001) and faster responses in the congruent condition (FAC: F_1,48_ = 34.116; p < 0.001) as compared to the neutral condition (Fig. 4). This was also reflected in a significant cumulative Stroop-effect (CUM: F_1,48_ = 215.247; p < 0.001). Participants got faster with practice, resulting in RTs being lower during post-exposure than pre-exposure testing (main effect of time, IF: F_1,48_ = 6.958; p = 0.011; FAC: F_1,48_ = 40.710; p < 0.001; CUM: F_1,48_ = 10.318; p = 0.002). We did not find main effects of exposure in any of the conditions (IF: F_1,48_ = 1.038; p = 0.313; FAC: F_1,48_ = 9.72e-6; p = 0.998; CUM: F_1,48_ = 0.936; p = 0.338). We also failed to find any veridical exposure effect in terms of interactions throughout all of our analyses. Unfortunately, in the incongruent condition, Real RTs differed from Sham RTs before exposure, in the baseline period, which resulted in several spurious interactions. In Supplementary Material 2 (Stroop Test Quality Control), we provide a detailed analysis of the Stroop test results (also involving the WN task) from the Sham condition.

**Figure 4.**
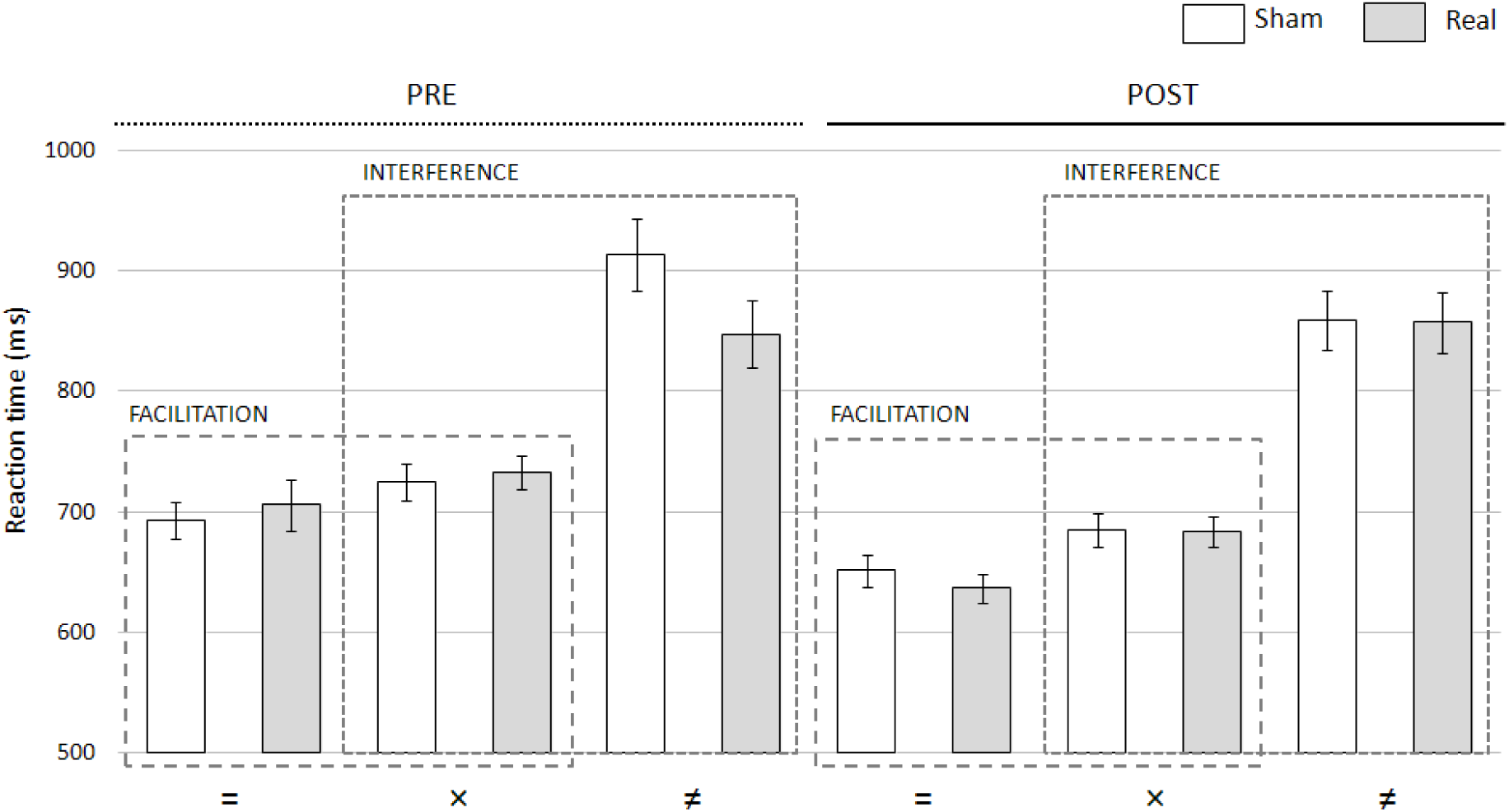
Comparison of RTs in the CN task pooled across RF types in the case of Facilitation and Interference before (Pre) and after (Post) RF exposure. Both FAC and IF appear both before and after exposure, but RF has no effect on either of the measures. Legends: [=] congruent, [×] neutral, [≠] incongruent. Error bars: Standard error of mean.

### EEG results

Alpha power significantly decreased in the RF exposure conditions relative to Sham control (main effect of exposure: F_1,53_ = 6.338; p = 0.015). Importantly, this effect was the result of differing temporal trends across exposure periods, as evidenced by a main effect of time (F_1.7,87.4_ = 7.309; p = 0.002) and a time × exposure interaction (F_1.8,95.3_ = 4.459; p = 0.017). A follow-up analysis on mid-exposure and post-exposure data with pre-exposure baseline values subtracted showed that the time × exposure interaction was driven by the exposure effect being stronger in the mid- and post-exposure periods compared to the pre-period (exposure in mid and post vs. baseline: F_1,53_ = 6.475; p = 0.014). However, the effect did not appear to differ between the mid-exposure and post-exposure periods (exposure × time mid vs. post: F_1,53_ = 2.202; p = 0.144).

We also performed the latter analysis (on baseline corrected data) on UMTS and LTE exposure separately. The results were analogous: alpha power significantly decreased as a result of LTE exposure (LTE exposure in mid and post vs. baseline: F_1,20_ = 5.095; p = 0.035), a similar (but marginally significant) effect was present for UMTS (UMTS exposure in Mid and Post vs. baseline: F_1,33_ = 3.772; p = 0.061), and the mid-post temporal interactions were not significant (exposure × time mid vs. post, UMTS: F_1,33_ = 0.037; p = 0.849; LTE: F_1,20_ = 2.932; p = 0.102). Also in line with this, analysis on the pooled data with RF type (UMTS vs. LTE) as a between-subject factor found no significant main effect of (RF type F_1,53_ = 0.724, p = 0.399) or interaction with RF type (time × RF type F_1,53_ = 2.927; p = 0.093; exposure × RF type F_1,53_ = 0.061; p = 0.806; time × exposure × RF type F_1,53_ = 1.562; p = 0.217).

The mass univariate analysis also yielded concordant results (Fig. 5). In the data pooled across RF types, a significant exposure effect was found (p_Cluster_ = 0.005). The topographic distribution of the difference covered the whole scalp in the mid- and post-exposure period (with only very subtle apparent differences between them), but with no difference in the pre-exposure period (top row of Fig. 5A). UMTS and LTE data analysed separately resulted in marginally significant differences with this method as well (UMTS: p_Cluster_ = 0.067, LTE: p_Cluster_ = 0.055; see Fig. 5A, second row, left and right topoplots for each time period), whereas testing for an RF type × exposure interaction yielded no suprathreshold clusters (see Fig. 5A, second row, smaller topoplots in the middle for each time period). Note that there are several apparent subtle differences in topographic patterns depending on exposure type and between the mid-exposure and post-exposure time periods, but as detailed above, these differences did not result in any significant interaction.

**Figure 5.**
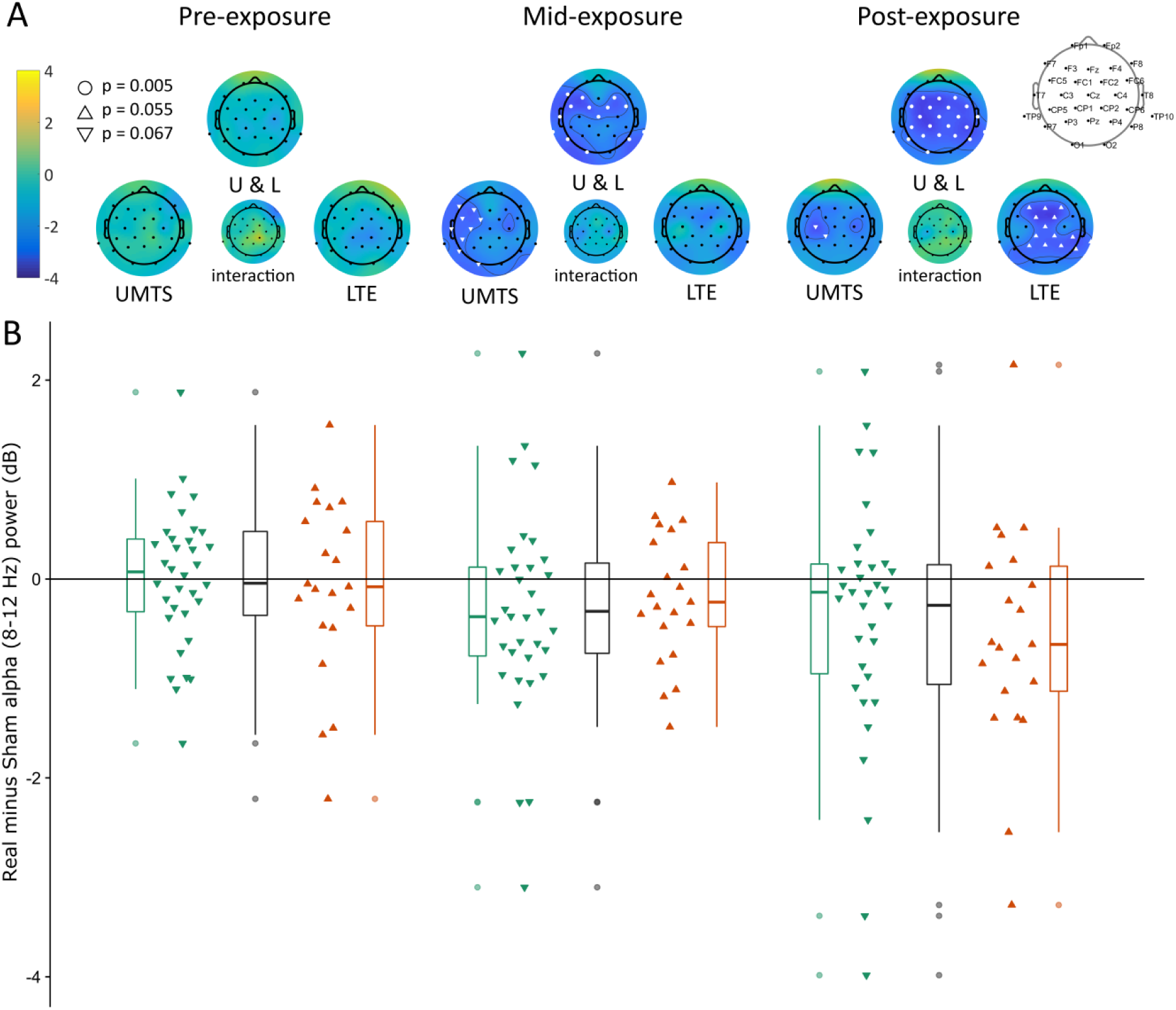
Real-Sham alpha band (8–12 Hz) power (dB) changes over the three exposure periods (pre-exposure, mid-exposure, post-exposure). (A) Topographic distribution of Real-Sham alpha band power difference. The colour scale bar represents t-values (left), white markers show suprathreshold clusters (cluster-corrected p-values for each cluster are shown in the legend on the left side). (B) Distributions of Real minus Sham alpha band power values. The layout matches that of panel A, boxplots and individual difference plots correspond to the large topographic plots (UMTS, UMTS and LTE together, LTE). On individual difference plots, one triangle marks the Real minus Sham difference in whole-scalp average alpha power of one subject in the corresponding time interval. Legends of boxes: line within the box: median (Q2); upper box edge: third quartile (Q3); lower box edge: first quartile (Q1); error bar upwards: Q3 + 1.5 × interquartile range (IQR) or maximum value, whichever is lower (IQR = Q3 – Q1); error bar downwards: Q1 - 1.5 × IQR or minimum value, whichever is greater; points: individual data points outside the range defined by the error bar (if any).

Our analyses focused on the alpha band that was a priori selected based on previous results. We do not present statistics from any other frequency range, but it is nevertheless important to verify that the spectral pattern of the results matches our expectations.^55^ In Fig. 6, it is evident that the effect is indeed confined to modulations of the peak in the alpha frequency band.

**Figure 6.**
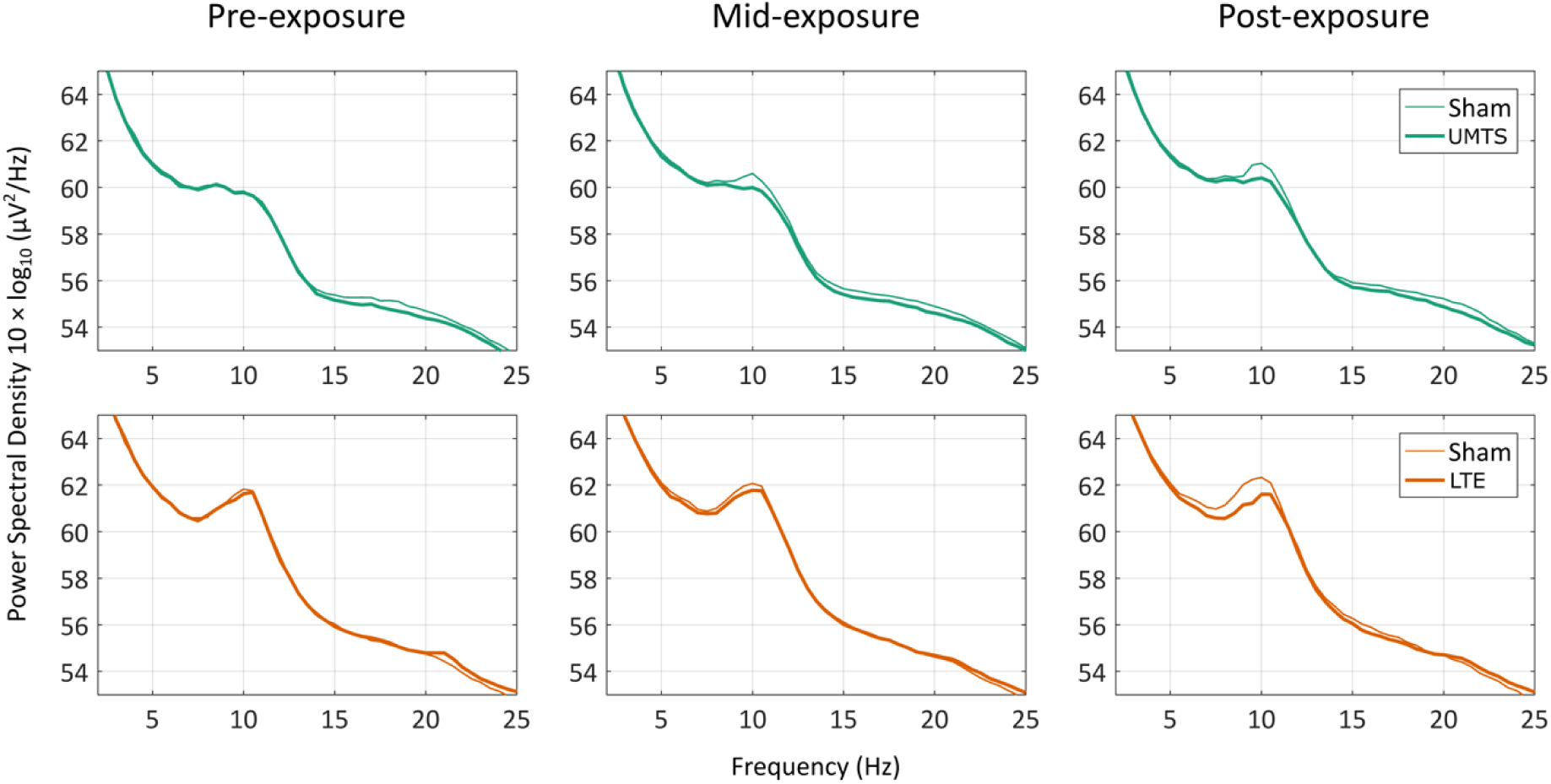
Pre-exposure, mid-exposure and post-exposure power spectral density from Real and Sham sessions (thick and thin lines, respectively) averaged over all EEG electrodes and across subjects in the UMTS and the LTE exposure groups.

With regard to our results of control analyses, neither the order of the Real and Sham exposure sessions (session order: F_1,53_ = 2.42; p = 0.13, time × session order: F_2,106_ = 1.143; p = 0.67, exposure × session order: F_1,53_ = 0.0877; p = 0.77, time × exposure × session order: F_2,106_ = 1.74; p = 0.18) nor the film order factors (film order: F_1,53_ = 0.135; p = 0.71, time × film order: F_2,106_ = 0.096; p = 0.91, exposure × film order: F_1,53_ = 0.43; p = 0.51, time × exposure × film order: F_2,106_ = 1.92; p = 0.15) displayed effects or interactions with any of the examined variables. Thus, we can exclude that these factors could have influenced our RF exposure related results.

## Discussion

We examined the effect of 3G and 4G RF EMF exposure on cognitive functions manifested in Stroop test performance and resting state spontaneous EEG in a double-blind, randomised, counterbalanced, crossover study.

UMTS and LTE exposure caused no measurable effects on the observed parameters of attentional processing in the Stroop test—neither on IF nor on FAC. Both IF and FAC were reliably present during the Real RF exposure in both experiments. Moreover, we did not find any change in the effect of RF even when we considered a more robust variable—the difference between RTs in congruent and incongruent situations. In contrary to the present results obtained in the acute exposure conditions, some epidemiological studies showed shorter^19^ or longer^16,17^ IF RTs in groups with higher frequency of MP use. It is possible that the change in performance found in the epidemiological studies was not directly caused by RF exposure itself but by other (confounding) factors. Evaluating long-term epidemiological studies (involving data about cognitive abilities, personality and user habits) together with well-controlled laboratory studies could help us to differentiate the general effects of MP using habits from the effects of the RF energy exposure itself.

Analysing the effects RF exposure on resting EEG with data pooled across the UMTS and LTE RF types revealed that, relative to the same time periods in Sham exposure sessions, alpha band power decreased both during and after irradiation. There was no difference between Sham and Real sessions in the pre-exposure baseline, and the exposure effects (decrease of alpha power) in the mid-exposure and post-exposure periods were also significant when assessed relative to the baseline period, but did not differ from each other. The effect was not at all focal to the site of irradiation, but rather appeared to be global, suggesting a brain-wide modulation of alpha oscillations by the RF exposure. UMTS and LTE exposure had similar effects, which imply that either the biophysical mechanism that mediates the neural effect should be to some extent shared between the two RF types or that the differences are too subtle to detect with the current data and methodology.

It is consistent that the EMF can affect the alpha band of resting EEG,^56^ although the reason why alpha power reacts differently (increases/decreases) to the RF exposure between studies remains unclear. In terms of waking EEG, the effect of RF (considering GSM systems) has been predominantly shown to increase alpha band power,^57-61^ although some reductions of alpha band power have also been reported.^56,62-64^ In a single-blind study by D’Costa *et al.*,^62^ during a single 25-min Real GSM exposure, statistically significant decreases were found in EEG power at 8 Hz and 9 Hz in the occipital region, and at 7 Hz and 9 Hz in both the occipital end central recording sites compared to Sham exposure. Perentos *et al.*^64^ observed a suppression of the global alpha band (8–12.75 Hz) activity under a 20-min low-level Real GSM-like extremely low frequency (ranging from direct current to at least 40 kHz) exposure compared to Sham in the exposed hemisphere only. In their follow-up study,^63^ it was found that this decrease in alpha band power was present also both under pulse modulated and continuous GSM RF exposure. Another article^56^ reported a statistically significant decrease of the alpha band spectral power in the post-exposure session after GSM RF exposure. The most relevant factors that may explain the bidirectional alpha power changes across the different studies are the use of diverse methods, intensities or frequencies and different experimental protocols as discussed by Loughran *et al.*^65^ Individual variability is also one of the factors that can cause discrepancies between the different results concerning the alpha power.^65,66^

A recent investigation of LTE exposure on resting EEG^67^ modulation in the alpha power was also detected. The double-blind, counterbalanced experiment involving 25 male adult subjects used a standard formulation for LTE signals and found reduced alpha band power during and after LTE exposure as compared to the pre-exposure period. In another recent LTE study,^68^ the functional connectivity reflected in the neural synchronisation of EEG signals was assessed, and LTE exposure was found to modulate the synchronisation patterns of brain activation also in areas remote to the exposure—even in the contralateral hemisphere. This phenomenon was similar to a previous finding of the same group based on fMRI data^69^ where a 30-min LTE RF exposure modulated spontaneous low frequency fluctuations in temporal and frontomedial brain regions, a finding which is also in line with other studies.^57,70^

The strength of alpha oscillations measurable on the scalp is associated with local modulations of excitability related to sensory or congnitive processes within an area: lower alpha power is often related to more intensive processing and higher cortical excitability.^71-73^ From this perspective, our results are compatible with the findings of recent transcranial magnetic stimulation (TMS) experiments showing incresased motor cortical excitability following acute exposure to GSM-EMF.^74^ Alpha oscillations are also thought to form important communication channels providing the basic scaffold of brain-wide visual attentional and memory networks.^75-77^ The fact that the exposure was localised to the right temporal cortex in our experiment and the alpha power decrease appeared as a global modulation over the whole scalp implies that local biophysical effects of the exposure in the temporal cortex may have led to global network-level changes in the brain. For example, it could be speculated that a change in the local signal-to-noise ratio in a few nodes of a network might require the whole network to adapt by re-tuning the sensitivity of oscillatory communication channels represented by alpha oscillations. The notion that, in the present experiment, the alpha modulation persisted even after the cessation of the irradiation could be due to the local changes also persisting, but also possibly from the temporal dynamics of the underlying network-level adaptation mechanism. Using the Stroop test, we could not show that such network-level alteration would be followed by or manifested in any observable changes in cognitive function. It is possible, again, that a different task might be more sensitive to network-level changes that might be reflected by the alpha modulation, but the presence of an EEG effect and the absence of a cognitive effect together suggest that the effects on neural processing might be subtle and stay well within the adaptation range of the functional brain networks involved in successful performance in the Stroop test.

## Conclusions

We can conclude that 20 min 3G (UMTS) or 4G (LTE) MP emitted RF exposure below the limit proposed by the International Commission of Non-Ionizing Radiation Protection (ICNIRP) does not interfere with executive function measures, processing speed and selective attention as assessed by the Stroop test. In addition, both RF exposure types caused a notable decrease in EEG alpha power over the whole scalp that persisted even after cessation of the exposure. These results imply that the brain networks underlying global alpha oscillations might require minor reconfiguration to adapt to the acute local biophysical changes in the temporal cortex caused by focal RF exposure mimicking MP use.

## Author Contributions

ZV, AT, PJ and IH designed research; ZV performed experiments; GT and PJ designed the exposure system; ZV, AT and BK analysed data; ZV, BK, IH and GT wrote the paper. All authors read and approved the final manuscript.

## Acknowledgements

We thank Viktória Báló and Norbert Zentai for their valuable assistance during data collection and Norbert Zentai for his kind IT support. The project was supported by the Hungarian National Program in Brain Sciences (NPIBS, KTIA NAP 13-1-2013-0001) and EFOP-3.6.2-16-2017-00008 „The role of neuro-inflammation in neurodegeneration: from molecules to clinics” at the University of Pécs. We thank all members of WG #2 at EU-COST Action BM1309 (EMF-MED) for valuable discussions on the experimental design and data interpretation.

## References

1 Kenechi Okeleke, M. R., Xavier Pedros. The Mobile Economy 2017. (© GSMA Intelligence 2017, 2017).

2 Gandhi, O. P. Electromagnetic fields: human safety issues. Annual Review of Biomedical Engineering 4, 211–234, doi:10.1146/annurev.bioeng.4.020702.153447 (2002).

3 Hyland, G. J. Physics and biology of mobile telephony. Lancet 356, 1833–1836 (2000).

4 Foster, K. R. & Glaser, R. Thermal mechanisms of interaction of radiofrequency energy with biological systems with relevance to exposure guidelines. Health Physics 92, 609–620, doi:10.1097/01.HP.0000262572.64418.38 (2007).

5 No.193., W. F. S. WHO Fact Sheet No.193. Electromagnetic fields and public health: mobile phones. (2014).

6 Ndoumbe Mbonjo Mbonjo, H. et al. Generic UMTS test signal for RF bioelectromagnetic studies. Bioelectromagnetics 25, 415–425, doi:10.1002/bem.20007 (2004).

7 Kwon, M. S. & Hamalainen, H. Effects of mobile phone electromagnetic fields: critical evaluation of behavioral and neurophysiological studies. Bioelectromagnetics 32, 253–272, doi:10.1002/bem.20635 (2011).

8 Zhang, J., Sumich, A. & Wang, G. Y. Acute effects of radiofrequency electromagnetic field emitted by mobile phone on brain function. Bioelectromagnetics 38, 329–338, doi:10.1002/bem.22052 (2017).

9 Kleinlogel, H. et al. Effects of weak mobile phone - electromagnetic fields (GSM, UMTS) on event related potentials and cognitive functions. Bioelectromagnetics 29, 488–497, doi:10.1002/bem.20418 (2008).

10 Kleinlogel, H. et al. Effects of weak mobile phone - electromagnetic fields (GSM, UMTS) on well-being and resting EEG. Bioelectromagnetics 29, 479–487, doi:10.1002/bem.20419 (2008).

11 Leung, S. et al. Effects of 2G and 3G mobile phones on performance and electrophysiology in adolescents, young adults and older adults. Clinical Neurophysiology 122, 2203–2216, doi:10.1016/j.clinph.2011.04.006 (2011).

12 Sauter, C. et al. Effects of Exposure to Electromagnetic Fields Emitted by GSM 900 and WCDMA Mobile Phones on Cognitive Function in Young Male Subjects. Bioelectromagnetics 32, 179–190, doi:10.1002/bem.20623 (2011).

13 Unterlechner, M., Sauter, C., Schmid, G. & Zeitlhofer, J. No effect of an UMTS mobile phone-like electromagnetic field of 1.97 GHz on human attention and reaction time. Bioelectromagnetics 29, 145–153, doi:10.1002/bem.20374 (2008).

14 Trunk, A. et al. Effects of concurrent caffeine and mobile phone exposure on local target probability processing in the human brain. Scientific Reports 5, 14434, doi:10.1038/srep14434 (2015).

15 Schmid, G., Sauter, C., Stepansky, R., Lobentanz, I. S. & Zeitlhofer, J. No influence on selected parameters of human visual perception of 1970 MHz UMTS-like exposure. Bioelectromagnetics 26, 243–250, doi:10.1002/bem.20076 (2005).

16 Abramson, M. J. et al. Mobile telephone use is associated with changes in cognitive function in young adolescents. Bioelectromagnetics 30, 678–686, doi:10.1002/bem.20534 (2009).

17 Redmayne, M. et al. Use of mobile and cordless phones and cognition in Australian primary school children: a prospective cohort study. Environmental Health 15, 26, doi:10.1186/s12940-016-0116-1 (2016).

18 Mortazavi, S. A., Tavakkoli-Golpayegani, A., Haghani, M. & Mortazavi, S. M. Looking at the other side of the coin: the search for possible biopositive cognitive effects of the exposure to 900 MHz GSM mobile phone radiofrequency radiation. Journal of Environmental Health Science and Engineering 12, 75, doi:10.1186/2052-336X-12-75 (2014).

19 Arns, M., Van Luijtelaar, G., Sumich, A., Hamilton, R. & Gordon, E. Electroencephalographic, personality, and executive function measures associated with frequent mobile phone use. The International Journal of Neuroscience 117, 1341–1360, doi:10.1080/00207450600936882 (2007).

20 Edelstyn, N. & Oldershaw, A. The acute effects of exposure to the electromagnetic field emitted by mobile phones on human attention. Neuroreport 13, 119–121 (2002).

21 Koivisto, M. et al. Effects of 902 MHz electromagnetic field emitted by cellular telephones on response times in humans. Neuroreport 11, 413–415 (2000).

22 Curcio, G. Exposure to Mobile Phone-Emitted Electromagnetic Fields and Human Attention: No Evidence of a Causal Relationship. Frontiers in Public Health 6, 42, doi:10.3389/fpubh.2018.00042 (2018).

23 Loughran, S. P. et al. No increased sensitivity in brain activity of adolescents exposed to mobile phone-like emissions. Clinical Neurophysiology, doi:10.1016/j.clinph.2013.01.010 (2013).

24 Haarala, C. et al. 902 MHz mobile phone does not affect short term memory in humans. Bioelectromagnetics 25, 452–456, doi:10.1002/bem.20014 (2004).

25 Hamblin, D. L., Croft, R. J., Wood, A. W., Stough, C. & Spong, J. The sensitivity of human event-related potentials and reaction time to mobile phone emitted electromagnetic fields. Bioelectromagnetics 27, 265–273, doi:10.1002/bem.20209 (2006).

26 Trunk, A. et al. Lack of interaction between concurrent caffeine and mobile phone exposure on visual target detection: an ERP study. Pharmacology, Biochemistry, and Behavior 124, 412–420, doi:10.1016/j.pbb.2014.07.011 (2014).

27 Valentini, E., Ferrara, M., Presaghi, F., De Gennaro, L. & Curcio, G. Systematic review and meta-analysis of psychomotor effects of mobile phone electromagnetic fields. Occupational and Environmental Medicine 67, 708–716, doi:10.1136/oem.2009.047027 (2010).

28 Barth, A., Ponocny, I., Gnambs, T. & Winker, R. No effects of short-term exposure to mobile phone electromagnetic fields on human cognitive performance: a meta-analysis. Bioelectromagnetics 33, 159–165, doi:10.1002/bem.20697 (2012).

29 Valentini, E., Ferrara, M., Presaghi, F., De Gennaro, L. & Curcio, G. Republished review: systematic review and meta-analysis of psychomotor effects of mobile phone electromagnetic fields. Postgraduate Medical Journal 87, 643–651, doi:10.1136/pgmj.2009.047027rep (2011).

30 Stroop. Studies of interference in serial verbal reactions. Journal of Experimental Psychology, 18, 643–662. (1935).

31 Treisman, A. & Fearnley, S. The Stroop test: selective attention to colours and words. Nature 222, 437–439 (1969).

32 Hutchison, K. A., Balota, D. A. & Duchek, J. M. The utility of Stroop task switching as a marker for early-stage Alzheimer’s disease. Psychology and Aging 25, 545–559, doi:10.1037/a0018498 (2010).

33 Koss, E., Ober, B. A., Delis, D. C. & Friedland, R. P. The Stroop color-word test: indicator of dementia severity. The International Journal of Neuroscience 24, 53–61 (1984).

34 Henik, A. & Salo, R. Schizophrenia and the stroop effect. Behavioral and Cognitive Neuroscience Reviews 3, 42–59, doi:10.1177/1534582304263252 (2004).

35 McGrath, J., Scheldt, S., Welham, J. & Clair, A. Performance on tests sensitive to impaired executive ability in schizophrenia, mania and well controls: acute and subacute phases. Schizophrenia Research 26, 127–137 (1997).

36 Assef, E. C., Capovilla, A. G. & Capovilla, F. C. Computerized stroop test to assess selective attention in children with attention deficit hyperactivity disorder. The Spanish Journal of Psychology 10, 33–40 (2007).

37 Lopez-Villalobos, J. A. et al. [Usefulness of the Stroop test in attention deficit hyperactivity disorder]. Revista de Neurologia 50, 333–340 (2010).

38 MacLeod, C. M. Half a century of research on the Stroop effect: an integrative review. Psychological Bulletin 109, 163–203 (1991).

39 Nunez, P. L. & Srinivasan, R. Electric fields of the brain: the neurophysics of EEG. (Oxford University Press, 2006).

40 Buzsaki, G. Rhythms of the Brain. (Oxford University Press, 2006).

41 Cohen, M. X. Where Does EEG Come From and What Does It Mean? Trends in Neurosciences 40, 208–218, doi:10.1016/j.tins.2017.02.004 (2017).

42 Gjoneska, B., Markovska-Simoska, S., Hinrikus, H., Pop-Jordanova, N. & Pop-Jordanov, J. Brain Topography of EMF-Induced EEG-Changes in Restful Wakefulness: Tracing Current Effects, Targeting Future Prospects. Prilozi 36, 103–112, doi:10.1515/prilozi-2015-0085 (2015).

43 Mueller, S. T. & Piper, B. J. The Psychology Experiment Building Language (PEBL) and PEBL Test Battery. Journal of Neuroscience Methods 222, 250–259, doi:10.1016/j.jneumeth.2013.10.024 (2014).

44 Luce, R. D. Response times: Their Role in Inferring Elementary Mental Organization. (Oxford University Press, 1986).

45 Heathcote, A., Popiel, S. J. & Mewhort, D. J. K. Analysis of Response-Time Distributions - an Example Using the Stroop Task. Psychological Bulletin 109, 340–347, doi:10.1037/0033-2909.109.2.340 (1991).

46 Whelan, R. Effective Analysis of Reaction Time Data. The Psychological Record 58, 475–482 (2008).

47 Ratcliff, R. Methods for dealing with reaction time outliers. Psychological Bulletin 114, 510–532 (1993).

48 Vecsei, Z., Csatho, A., Thuroczy, G. & Hernadi, I. Effect of a single 30 min UMTS mobile phone-like exposure on the thermal pain threshold of young healthy volunteers. Bioelectromagnetics 34, 530–541, doi:10.1002/bem.21801 (2013).

49 Trunk, A. et al. No effects of a single 3G UMTS mobile phone exposure on spontaneous EEG activity, ERP correlates, and automatic deviance detection. Bioelectromagnetics 34, 31–42, doi:10.1002/bem.21740 (2013).

50 Parazzini, M. et al. Effects of UMTS cellular phones on human hearing: results of the European project EMFnEAR. Radiation Research 172, 244–251, doi:10.1667/RR1679.1 (2009).

51 Stefanics, G., Thuroczy, G., Kellenyi, L. & Hernadi, I. Effects of twenty-minute 3G mobile phone irradiation on event related potential components and early gamma synchronization in auditory oddball paradigm. Neuroscience 157, 453–462, doi:10.1016/j.neuroscience.2008.08.066 (2008).

52 Delorme, A. & Makeig, S. EEGLAB: an open source toolbox for analysis of single-trial EEG dynamics. Journal of Neuroscience Methods 134, 9–21 (2004).

53 Oostenveld, R., Fries, P., Maris, E. & Schoffelen, J. M. FieldTrip: Open source software for advanced analysis of MEG, EEG, and invasive electrophysiological data. Computational Intelligence and Neuroscience 2011, 156869, doi:10.1155/2011/156869 (2011).

54 Maris, E. & Oostenveld, R. Nonparametric statistical testing of EEG- and MEG-data. Journal of Neuroscience Methods 164, 177–190, doi:10.1016/j.jneumeth.2007.03.024 (2007).

55 van Ede, F. & Maris, E. Physiological Plausibility Can Increase Reproducibility in Cognitive Neuroscience. Trends in Cognitive Sciences 20, 567–569, doi:10.1016/j.tics.2016.05.006 (2016).

56 Ghosn, R. et al. Radiofrequency signal affects alpha band in resting electroencephalogram. Journal of Neurophysiology 113, 2753–2759, doi:10.1152/jn.00765.2014 (2015).

57 Croft, R. J. et al. Acute mobile phone operation affects neural function in humans. Clinical neurophysiology: official journal of the International Federation of Clinical Neurophysiology 113, 1623–1632 (2002).

58 Croft, R. J. et al. The effect of mobile phone electromagnetic fields on the alpha rhythm of human electroencephalogram. Bioelectromagnetics 29, 1–10, doi:10.1002/bem.20352 (2008).

59 Croft, R. J. et al. Effects of 2G and 3G mobile phones on human alpha rhythms: Resting EEG in adolescents, young adults, and the elderly. Bioelectromagnetics 31, 434–444, doi:10.1002/bem.20583 (2010).

60 Curcio, G. et al. Is the brain influenced by a phone call? An EEG study of resting wakefulness. Neuroscience Research 53, 265–270, doi:10.1016/j.neures.2005.07.003 (2005).

61 Regel, S. J. et al. Pulsed radio frequency radiation affects cognitive performance and the waking electroencephalogram. Neuroreport 18, 803–807, doi:10.1097/WNR.0b013e3280d9435e (2007).

62 D’Costa, H. et al. Human brain wave activity during exposure to radiofrequency field emissions from mobile phones. Australasian Physical & Engineering Sciences in Medicine 26, 162–167 (2003).

63 Perentos, N., Croft, R., McKenzie, R. & Cosic, I. The Alpha Band of the Resting Electroencephalogram under Pulsed and Continuous Radiofrequency Exposures. IEEE Transactions on Bio-Medical Engineering, 60, 1702–1710, doi:10.1109/TBME.2013.2241059 (2013).

64 Perentos, N., Croft, R. J., McKenzie, R. J., Cvetkovic, D. & Cosic, I. The effect of GSM-like ELF radiation on the alpha band of the human resting EEG. Conference proceedings: … Annual International Conference of the IEEE Engineering in Medicine and Biology Society. IEEE Engineering in Medicine and Biology Society. Conference 2008, 5680–5683, doi:10.1109/IEMBS.2008.4650503 (2008).

65 Loughran, S. P., McKenzie, R. J., Jackson, M. L., Howard, M. E. & Croft, R. J. Individual differences in the effects of mobile phone exposure on human sleep: rethinking the problem. Bioelectromagnetics 33, 86–93, doi:10.1002/bem.20691 (2012).

66 Hinrikus, H., Bachmann, M., Lass, J., Karai, D. & Tuulik, V. Effect of low frequency modulated microwave exposure on human EEG: individual sensitivity. Bioelectromagnetics 29, 527–538, doi:10.1002/bem.20415 (2008).

67 Yang, L., Chen, Q., Lv, B. & Wu, T. Long-Term Evolution Electromagnetic Fields Exposure Modulates the Resting State EEG on Alpha and Beta Bands. Clinical EEG and Neuroscience 48, 168–175, doi:10.1177/1550059416644887 (2017).

68 Lv, B., Su, C., Yang, L., Xie, Y. & Wu, T. Whole brain EEG synchronization likelihood modulated by long term evolution electromagnetic fields exposure. Conference proceedings: … Annual International Conference of the IEEE Engineering in Medicine and Biology Society. IEEE Engineering in Medicine and Biology Society. Conference 2014, 986–989, doi:10.1109/EMBC.2014.6943758 (2014).

69 Lv, B. et al. The alteration of spontaneous low frequency oscillations caused by acute electromagnetic fields exposure. Clinical Neurophysiology 125, 277–286, doi:10.1016/j.clinph.2013.07.018 (2014).

70 Haarala, C. et al. Effects of a 902 MHz mobile phone on cerebral blood flow in humans: a PET study. Neuroreport 14, 2019–2023, doi:10.1097/01.wnr.0000090954.15465.94 (2003).

71 Pfurtscheller, G. & Aranibar, A. Event-related cortical desynchronization detected by power measurements of scalp EEG. Electroencephalography and Clinical Neurophysiology 42, 817–826 (1977).

72 Foxe, J. J. & Snyder, A. C. The Role of Alpha-Band Brain Oscillations as a Sensory Suppression Mechanism during Selective Attention. Frontiers in Psychology 2, 154, doi:10.3389/fpsyg.2011.00154 (2011).

73 Jensen, O. & Mazaheri, A. Shaping functional architecture by oscillatory alpha activity: gating by inhibition. Frontiers in Human Neuroscience 4, 186, doi:10.3389/fnhum.2010.00186 (2010).

74 Ferreri, F. et al. Mobile phone emissions and human brain excitability. Annals of Neurology 60, 188–196, doi:10.1002/ana.20906 (2006).

75 Palva, S. & Palva, J. M. New vistas for alpha-frequency band oscillations. Trends in Neurosciences 30, 150–158, doi:10.1016/j.tins.2007.02.001 (2007).

76 Michalareas, G. et al. Alpha-Beta and Gamma Rhythms Subserve Feedback and Feedforward Influences among Human Visual Cortical Areas. Neuron 89, 384–397, doi:10.1016/j.neuron.2015.12.018 (2016).

77 Fries, P. Rhythms for Cognition: Communication through Coherence. Neuron 88, 220–235, doi:10.1016/j.neuron.2015.09.034 (2015).

